# Bumblebee occupancy responds to complex interactions between local and landscape land use, climatic niche properties and climate change

**DOI:** 10.1101/2023.09.12.557199

**Authors:** Tim Newbold, Jeremy Kerr, Peter Soroye, Jessica J. Williams

**Affiliations:** Centre for Biodiversity and Environment Research, Department of Genetics, Evolution and Environment, University College London, London, UK; Department of Biology, University of Ottawa, Ottawa, Ontario, Canada

## Abstract

Insect biodiversity is changing rapidly, driven by a complex suite of pressures, foremost among which are human land use, land-use intensification, and increasingly climate change. Bumblebees deliver important pollination services to wild plants and human crops, but we lack large-scale empirical evidence on how land use and climate change interact to drive bumblebee biodiversity changes. We assess bumblebee occupancy responses to interactive effects of land use and climate pressures across North America and Western Europe. Occupancy increases with landscape natural habitat and decreases with the duration of human use of landscapes. Responses to historical climate warming are negative in natural habitats but positive in human land uses, while human land use reduces occupancy most in the centre of species’ temperature niches. We estimate that the combined pressures have reduced bumblebee occupancy by 61% across sampled natural habitats, and 65% across human land uses, suggesting that treating present-day natural habitats as an undisturbed reference is misleading. Our results can inform efforts to conserve bumblebee biodiversity in the face of ongoing land-use changes and accelerating climatic changes.

**One-sentence summary:** Land use and climate change interact to drive large declines in bumblebee occupancy in both natural and human-modified habitats

## Introduction

Insect biodiversity has been undergoing rapid changes in recent decades, although reported trends differ in direction and magnitude. Several studies have suggested steep declines in the abundance, richness and distributions of terrestrial insect species, including bumblebees (*1–5*). On the other hand, other studies have suggested a more mixed picture, with little overall change in insect biodiversity on average (*6*, *7*). Biodiversity changes, including those of insects, are often characterized by strong turnover, with certain species - or groups of species - showing more positive trends than others (*6*, *8–10*).

Human land use and land-use intensification have been identified as key drivers of insect biodiversity changes. In general, previous studies have demonstrated a negative effect on bees (including bumblebees) of the conversion of natural habitats to agriculture and other human-modified habitats (*5*, *11–13*), although responses vary strongly among species (*14*, *15*) and among regions (*16*). Furthermore, within agricultural areas, intensification of farming practices (e.g. lowering of crop diversity, or the removal of flower-rich field margins) is associated with further reductions, on average, in bee diversity (*17*, *18*). One facet of agricultural intensification that is of particular importance for bees is the increased application of chemical pesticides, which has been associated with reductions in bee distributions, population persistence, and reproductive performance (*19–23*), although effects vary spatially (*24*).

The effects of land use and habitat differences can operate on larger spatial and temporal scales than captured at the location and time of biodiversity sampling. The availability of natural or semi-natural habitats in the landscape has been shown to be important for sustaining the diversity of bees, as well as improving individual and colony performance, including in agricultural habitats (*12*, *25–28*). However, effects often depended on the species and/or the nature of the natural habitat (*8*, *28*, *29*), and sometimes were absent or negative, for example when farmed areas provide abundant floral resources (*25*, *30*). At the same time, farming practices within agricultural land uses may impact biodiversity within natural habitats. For example, pesticides and other agricultural chemicals may disperse out of the farmland in which they are applied (*31*), and so could impact insect biodiversity across all habitats in a landscape. Similarly, over time, agricultural landscapes tend to become progressively intensified, becoming more homogeneous and losing remnant natural habitats (*32*). Thus, we might expect landscapes that have been farmed for longer to contain a lower diversity of bees and other insects.

Climate change is already exerting a strong influence on patterns of bee and bumblebee biodiversity, and its effect is likely to grow rapidly in the coming years. Already, bumblebee distributions have been contracting at the southern range margin (*1*) and moving to higher elevations (*33*), while the Community Temperature Index of bumblebee communities (which measures the average temperature affiliation of species in a community) has been increasing (*34*), consistent with expectations under climate warming. Indeed, analyses of changes in bee diversity over time have revealed an important role for warming temperatures in observed declines (*3*, *5*, *8*), and have revealed larger declines in more climatically specialized species (*35*). Experimental exposure of bumblebees to heat stress led to high rates of mortality in bumblebee species (*36*). Future projections suggest that the impact of climate change on bumblebees is likely to increase in the coming decades (*14*, *37*).

Recent evidence points toward synergistic interactive effects of land use, land-use intensification and climate change on biodiversity, but only a small number of studies have focused on insects (*38–41*). These interactive effects are driven by two key mechanisms. First, habitat disturbance impedes species shifting their ranges in response to climate change (*42*). Second, conversion of natural habitats to agriculture and other human uses changes local climates, typically towards hotter and drier conditions with greater extremes of temperature (*42*, *43*). For insects in general, the coincidence of intensive agricultural land use and rapid recent climate change is associated with reductions of around 50% of total assemblage abundance, with the greatest losses occurring where little natural habitat remains in the landscape (*41*). Insect species tolerant of warmer and drier conditions, on average, tend to be favoured in agricultural and urban land uses (*38–40*), probably as a result of the altered local climatic conditions in these areas. Studies on vertebrates have revealed that populations living close to the upper realized temperature niche of species respond more negatively to human land uses than populations living elsewhere (*44*).

Bumblebees are a key group of species, of high importance for both natural ecosystems and agricultural production. Half of plant species are estimated to depend on wild animal pollinators for more than 80% of their seed production (*45*). Lower pollinator biodiversity has been shown to reduce the reproductive success of wild plants (*46*), and greater rates of declines of plants that depend on animal pollination than other plant species may be associated with declines in key pollinator species (*47*). Many studies have shown positive relationships between pollinator diversity and measures of crop success (*48–52*). As ‘buzz’ pollinators (*53*), which use vibrations to dislodge pollen from flowers and play a vital role in the pollination of certain plant species, bumblebees make a substantial contribution to the pollination of many important crops, including apples, pears, melons, peppers, cucumbers, tomatoes, and many nuts and soft fruits (*54*).

Here we present a continental-scale study that simultaneously considers the interacting effects of climate change, land use and land-use intensification on bumblebee species’ occupancy. Despite there being numerous studies on the impact of individual pressures on bees, bumblebees and other insects, continental-scale studies remain rare, which has prevented the identification of general patterns in bee responses to interacting large-scale pressures. We study the impact of land-use-climate interactions on bumblebees across the continents of North America and Europe (although the available data were biased towards Europe). Specifically, we test the response of bumblebee occupancy to five classes of environmental variables: land use (natural habitat versus agricultural areas), land-use intensity (application rates of pesticides and fertilizer), land-use history, landscape natural habitat availability, and climate-related variables (population position within species’ realized temperature niche, and change in this niche position caused by recent climate change). We hypothesize that bumblebee occupancy will be lower in agricultural areas, particularly where land-use intensity is higher, where landscapes have been dominated by human activities for longer, and where less natural habitat remains within the landscape. We further hypothesize that occupancy will be most strongly reduced in agricultural land use for populations near the upper limit of species’ temperature niche, and especially where there have been recent increases in temperature. We test these hypotheses using mixed-effects models that fit bumblebee occupancy at sites across North America and Western Europe as a function of variables describing land use, land-use intensity and climatic niche properties, using data from the PREDICTS database (*55*, *56*). We include random effects to control for variation among the individual studies collated within the PREDICTS database, as well as random variation in occupancy among sites and species. Because the PREDICTS database contains snapshot spatial comparisons of biodiversity among different habitats, mostly sampled between 2000 and 2013, biodiversity may already have been reduced within natural habitats as a result of historic human pressures (leading to a ‘shifting baseline syndrome’ (*57*)). To test for such impacts, we use our models, which describe responses to multiple interacting pressures, to infer how bumblebee occupancy has been altered across sampled habitats, both natural and human-modified.

## Results

Availability of natural habitat in the landscape, land-use intensity, land-use history, climatic niche properties and climate change all made a substantial contribution to explaining bumblebee occupancy across land uses (Table 1). A model containing just land use was only marginally better than the null model (ΔAIC = -1), and land use alone explained only a very small proportion of the variation in bumblebee occupancy (R^2^_marginal_ = 0.0008; R^2^_conditional_ = 0.86; Table 1). The most complex, and best-fitting model, containing all of the candidate fixed effects, showed a much better fit than the null model (ΔAIC = -160.45), and explained substantially more of the variation in bumblebee occupancy (R^2^_marginal_ = 0.066; R^2^_conditional_ = 0.89; Table 1).

**Table 1:**
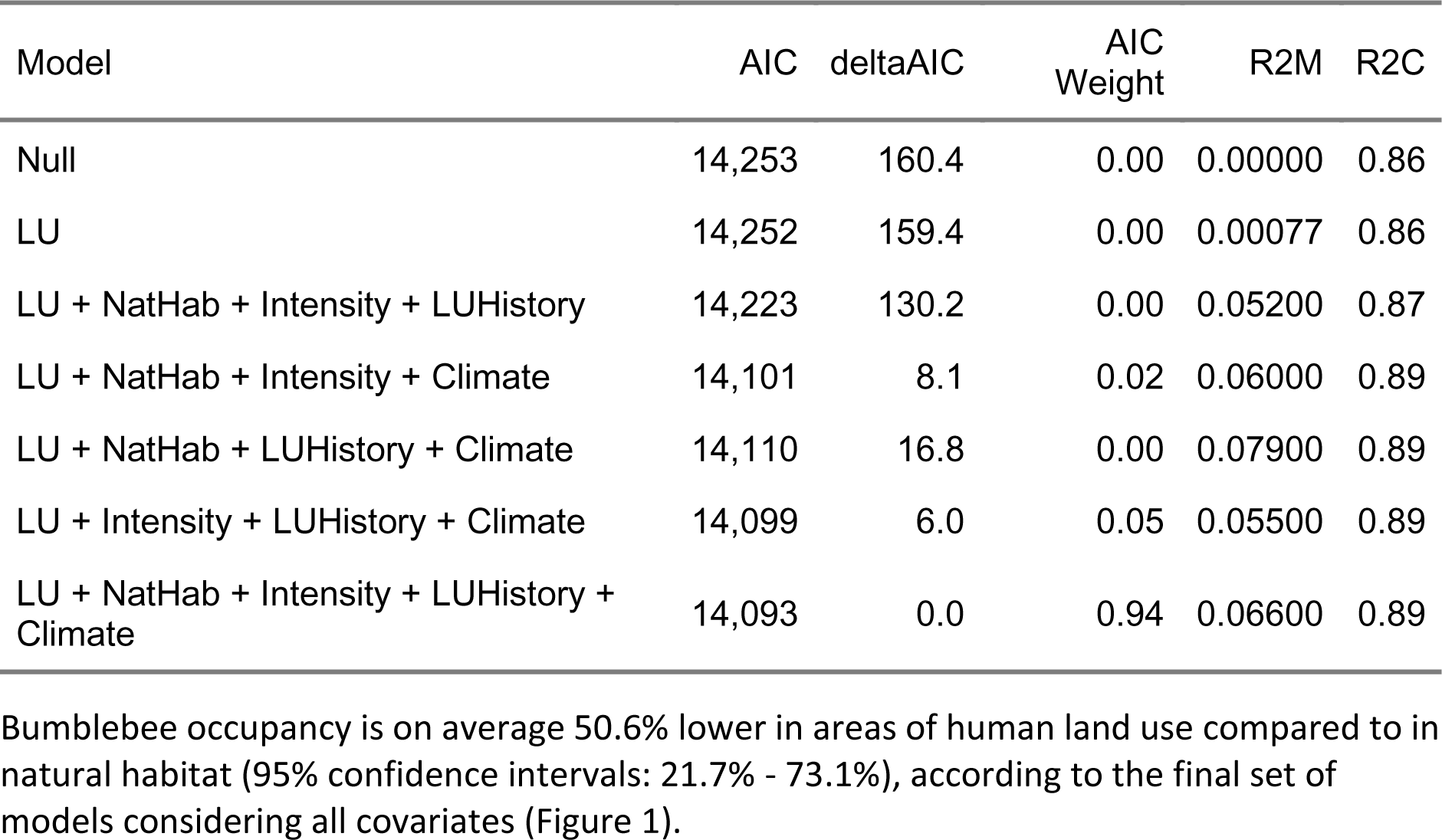
Summary statistics of the candidate models. Candidate models considered different combinations of variable categories: land use (LU), amount of natural habitat in the surrounding landscape (NatHab), land-use intensity variables (Intensity, consisting of rates of pesticide and fertilizer application), landscape history (LUHistory, specifically the duration of time since the landscape was substantially modified by human land uses) and climate-related variables (Climate, consisting of the realized temperature niche position in the baseline period (1901-1975) and the change in niche position between this baseline period and a more recent period (2000-2014). Statistics shown are: model AIC, the difference in AIC between the best model and the candidate model (deltaAIC), the AIC weight of the candidate model (Weight), and the marginal and conditional pseudo-R^2^ values of the candidate model (R2M and R2C, respectively)

| Model | AIC | deltaAIC | AIC Weight | $R^2_M$ | $R^2_C$ |
| --- | --- | --- | --- | --- | --- |
| Null | 14,253 | 160.4 | 0.00 | 0.00000 | 0.86 |
| LU | 14,252 | 159.4 | 0.00 | 0.00077 | 0.86 |
| LU + NatHab + Intensity + LUHistory | 14,223 | 130.2 | 0.00 | 0.05200 | 0.87 |
| LU + NatHab + Intensity + Climate | 14,101 | 8.1 | 0.02 | 0.06000 | 0.89 |
| LU + NatHab + LUHistory + Climate | 14,110 | 16.8 | 0.00 | 0.07900 | 0.89 |
| LU + Intensity + LUHistory + Climate | 14,099 | 6.0 | 0.05 | 0.05500 | 0.89 |
| LU + NatHab + Intensity + LUHistory + Climate | 14,093 | 0.0 | 0.94 | 0.06600 | 0.89 |

The presence or absence of bumblebee species in both natural habitats and areas of human-modified land use is shaped strongly by habitat characteristics of the surrounding landscape. In both land-use types, bumblebees are more likely to occur in landscapes with more natural habitat and that have been dominated by humans for a shorter length of time, with the effect of natural habitat strongest in areas of human land use (Figure 2).

**Figure 1:**
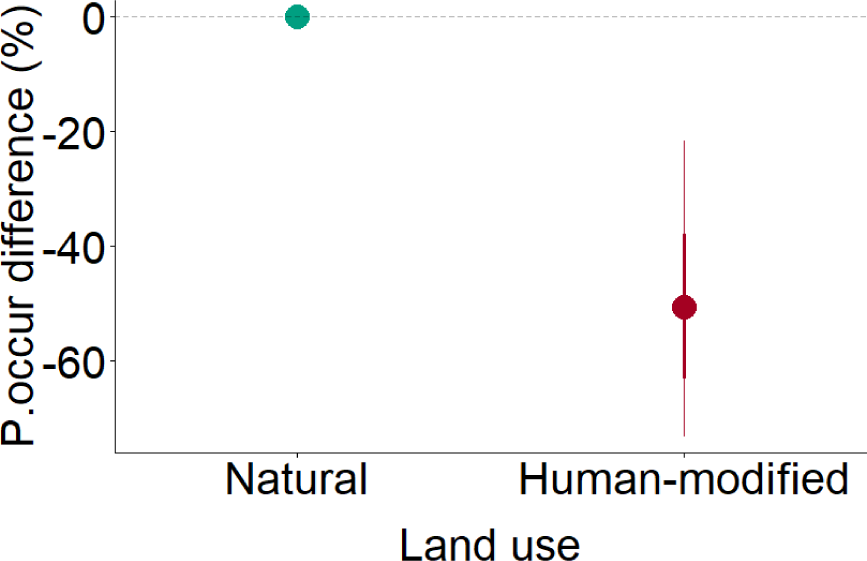
Effect of land use on bumblebee probability of occurrence. Values for human-modified habitat (i.e., agricultural and urban areas) are expressed as the % difference in probability of occurrence compared to natural (primary and secondary) habitats. All other variables are held at median values for the respective land use, except for baseline temperature niche position that is held at a fixed median value across both land-use types, and the change in temperature niche position owing to recent climate change that is held at zero. Points show the median prediction, while thick and thin error bars, respectively, represent the 67% and 95% confidence intervals. Predictions capture both model and parameter uncertainty, by drawing 1000 sets of estimates both across models (weighted by AIC weight), and within sampled models according to the modelled uncertainty in the parameter estimates

**Figure 2:**
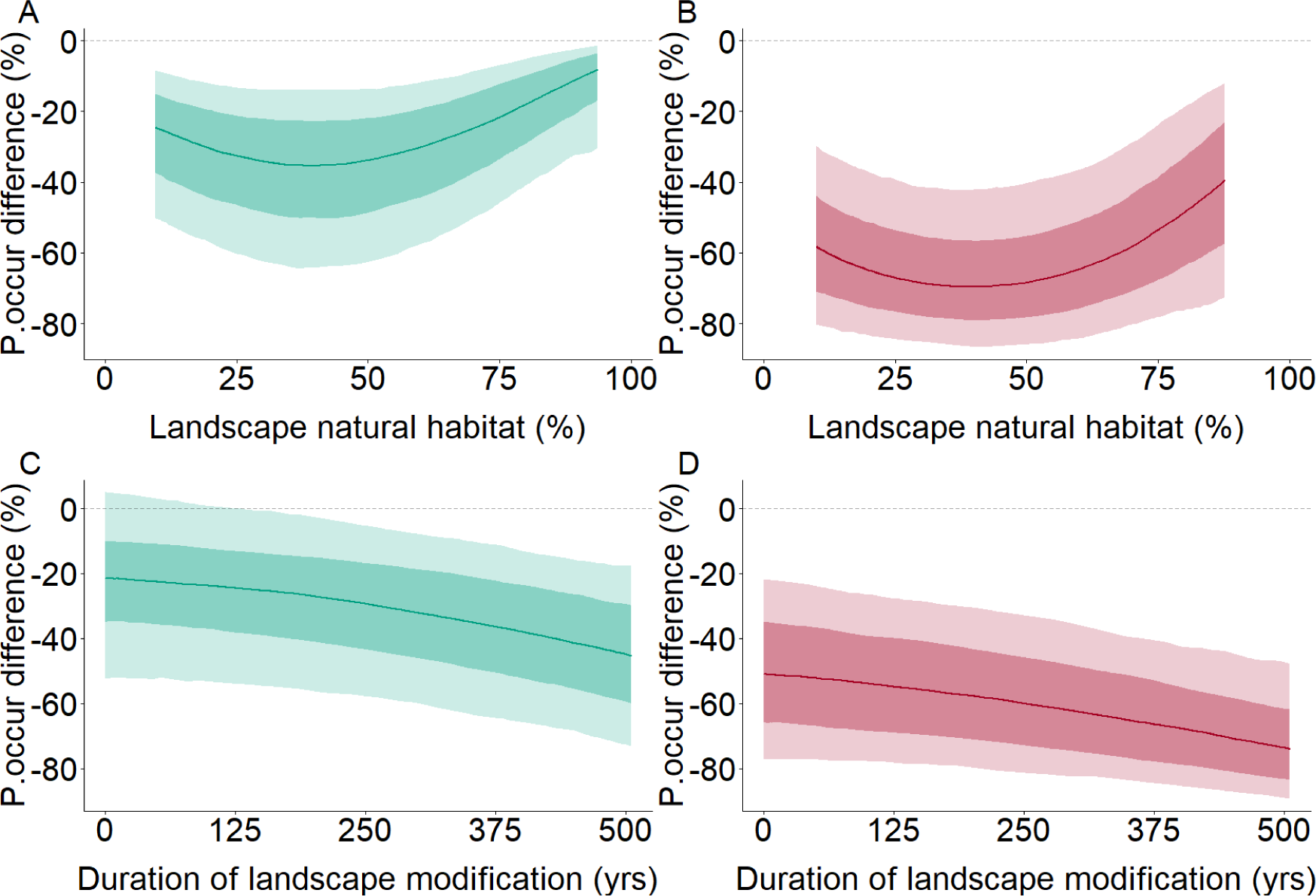
Effect of landscape-level land-use characteristics on bumblebee probability of occurrence across land uses. Landscape characteristics considered were the percentage of natural habitat (A and B) and the duration of substantial human modification of the landscape (i.e., the number of years since 30% of the landscape was converted to human-modified habitats) (C and D). Modelled effects are shown for natural (primary and secondary) habitats (A and C) and for human-modified (agricultural and urban) habitats (B and D). All other variables are held at median values for the respective land use, except for baseline temperature niche position that is held at a fixed value across both land-use types of 0.59 (the 10th percentile of values in the original dataset, representing a position near the temperature niche centre), and the change in temperature niche position owing to recent climate change that is held at zero. Values are expressed as the % difference in probability of occurrence compared to natural habitat locally, in a landscape composed entirely of natural habitat, with no history of landscape modification, with no application of agricultural chemicals, and no recent climate change. Lines represent modelled median projections, while dark and light shading, respectively, represent the 67% and 95% confidence intervals. Predictions capture both model and parameter uncertainty, by drawing 1000 sets of estimates both across models (weighted by AIC weight), and within sampled models according to the modelled uncertainty in the parameter estimates. Relationships are plotted for the central 95% of values sampled within each land use

Bumblebee occupancy is also shaped by rates of chemical input to the landscape, although less strongly than by landscape habitat characteristics. In natural habitats, the probability of occurrence of bumblebees decreases with increasing pesticide application, and shows a u-shaped relationship with fertiliser application (Figure 3A,C). In areas of human land use, probability of occurrence increases weakly with both pesticide and fertiliser application, with the latter relationship also being u-shaped (Figure 3B,D). Modelled effects of pesticide application rate were very similar when using either the low or high estimates of rates from the original dataset (*58*) (results not shown).

**Figure 3:**
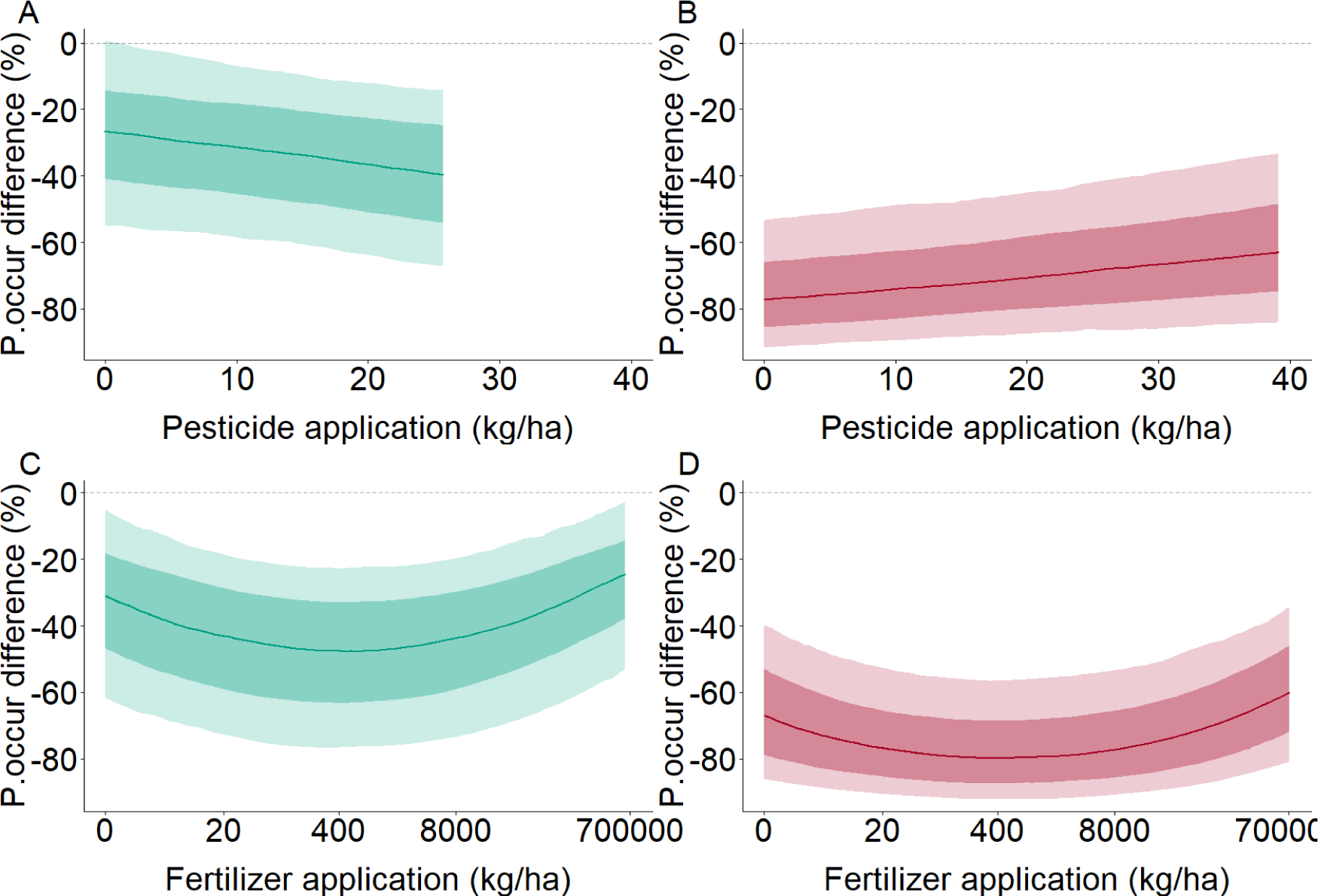
Effect of chemical inputs on bumblebee probability of occurrence across land uses. We considered the density of application of pesticides (A and B) and fertilizers (C and D). Modelled effects are shown for natural (primary and secondary) habitats (A and C) and for human-modified (agricultural and urban) habitats (B and D). All other variables are held at median values for the respective land use, except for baseline temperature niche position that is held at a fixed value across both land-use types of 0.59 (the 10th percentile of values in the original dataset, representing a position near the temperature niche centre), and the change in temperature niche position owing to recent climate change that is held at zero. Values are expressed as the % difference in probability of occurrence compared to natural habitat locally, in a landscape composed entirely of natural habitat, with no history of landscape modification, with no application of agricultural chemicals, and no recent climate change. Lines represent modelled median projections, while dark and light shading, respectively, represent the 67% and 95% confidence intervals. Predictions capture both model and parameter uncertainty, by drawing 1000 sets of estimates both across models (weighted by AIC weight), and within sampled models according to the modelled uncertainty in the parameter estimates. Relationships are plotted for the central 95% of values sampled within each land use

Finally, bumblebee probability of occurrence is strongly influenced by the climatic niche properties of species, and how these properties have been modified by recent climate change. Ignoring the effects of climate change initially, populations near the centre of the species’ temperature niche showed the greatest declines in probability of occurrence between natural and human-modified habitats, whereas for populations nearer the upper limit of the temperature niche, there was little difference in occupancy (Figure 4A,B). An increase in realized temperature niche position caused by climate warming had a negative effect on probability of occurrence in natural habitats, especially for populations near the species’ temperature niche centre (Figure 4C). In contrast, the effect of climate warming in human land uses was less clear, and there was a weak positive effect if anything (Figure 4D).

**Figure 4:**
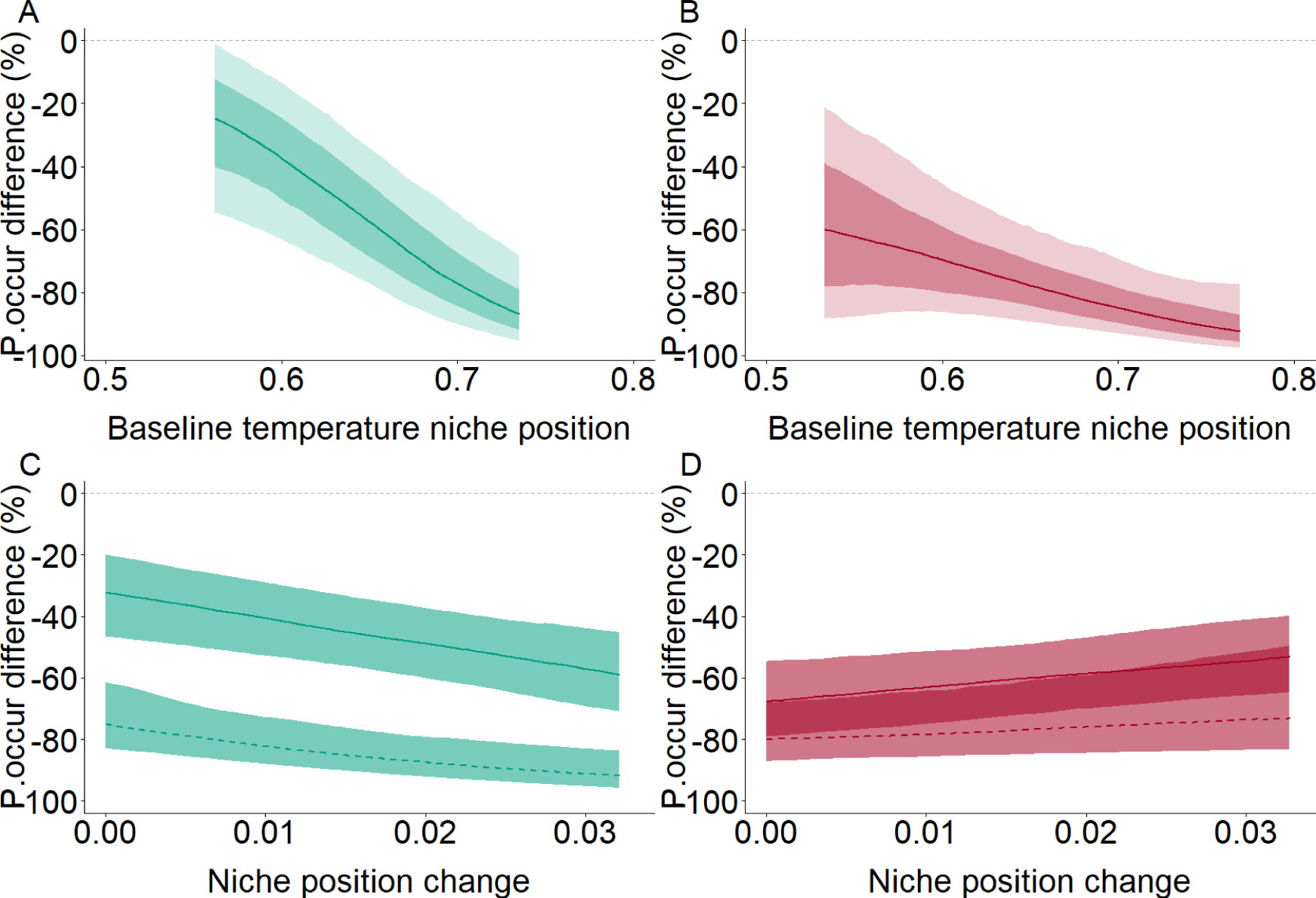
Effect of climatic niche properties and climate change on bumblebee probability of occurrence across land uses. We considered the effects of realized temperature niche position in a baseline period (1901 - 1975) (A and B) and change in realized temperature niche position between the baseline and a recent period (2000 - 2014) (C and D). A temperature niche position value of 0 represents a population at the cold edge of the species’ realized temperature niche, while a value of 1 represents a population at the warm edge of the species’ realized temperature niche. Modelled effects are shown for natural (primary and secondary) habitats (A and C) and for human-modified (agricultural and urban) habitats (B and D). For change in temperature niche position, we further sub-divide modelled responses according to baseline temperature niche position, plotting modelled relationships for populations near the species’ temperature niche centre (a value of 0.59, representing the 10th percentile of values sampled in the original dataset; solid lines) and species nearer the niche edge (a value of 0.73, representing the 90th percentile of values sampled in the original dataset; dashed lines). All other variables are held at median values for the respective land use. Values are expressed as the % difference in probability of occurrence compared to natural habitat locally, in a landscape composed entirely of natural habitat, with no history of landscape modification, with no application of agricultural chemicals, and no recent climate change. Lines represent modelled median projections, while dark and light shading, respectively, represent the 67% and 95% confidence intervals. Predictions capture both model and parameter uncertainty, by drawing 1000 sets of estimates both across models (weighted by AIC weight), and within sampled models according to the modelled uncertainty in the parameter estimates. Relationships are plotted for the central 95% of values sampled within each land use

The models deviated slightly from the assumptions of standard parametric statistical tests (Figure S1). Nevertheless, very similar model-predicted values were obtained when refitting the final best-fitting model using MCMC, which is a fundamentally different approach to fitting estimates of model parameters (Figure S2). Spatial autocorrelation was detected in the residuals associated with a slightly higher fraction (11.8%) of underlying studies than would be expected by chance (Figure S3).

Using the models to infer bumblebee probability of occupancy at the sampled sites, compared to the probability under hypothetical baseline conditions (natural habitat locally and across the whole landscape, no history of human land use, no application of agricultural chemicals, and no climate change), we estimate that bumblebee occupancy has been reduced by 58% (95% CI: 33% – 77%) across sampled natural habitats, and by 65% (95% CI: 36% – 82%) across sampled human land uses (Figure 5).

**Figure 5:**
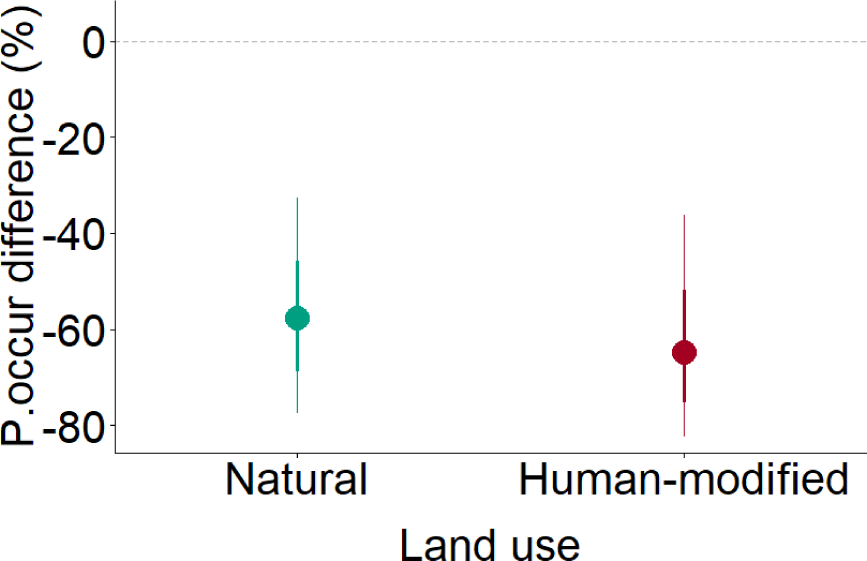
Model-inferred differences in average bumblebee probability of occupancy at sampled sites in natural (primary and secondary) habitats and human-modified (agricultural and urban) habitats, compared to hypothetical baseline conditions (natural land use across the whole landscape, no human land-use history, no agricultural chemical application, and no climate change). Predictions were generated for every species recorded at each sampled site across both natural and human-modified habitats, based on the values of the explanatory variables as estimated for the sampled sites. We then average the predictions across all species and all sampled sites to generate the average predictions for natural and human-modified habitats. Points show the median prediction, while thick and thin error bars, respectively, represent the 67% and 95% confidence intervals. Predictions capture both model and parameter uncertainty, by drawing 1000 sets of estimates both across models (weighted by AIC weight), and within sampled models according to the modelled uncertainty in the parameter estimates

## Discussion

Our results reveal important interactive effects of climate change and land use on bumblebee biodiversity. There is a growing recognition that synergistic interactions among pressures, especially land-use-climate interactions, may be exacerbating ongoing biodiversity changes (*42*, *43*), but relatively few studies have investigated such interactions in the context of insect biodiversity change (*38–41*). While a correlative empirical analysis such as ours cannot confidently elucidate mechanisms, it is likely that two mechanisms play a key role: microclimatic changes associated with land-use change; and habitat fragmentation impeding responses to climate change (*42*). We estimate that bumblebee occupancy has been reduced substantially across both natural habitats and human-modified areas. As climate changes intensify, and further modification of land use occurs, changes in bumblebee biodiversity are likely to accelerate.

We find that bumblebee occupancy is reduced most in areas of human land use, especially in landscapes with a lower cover of natural habitats that have been modified by humans for a long period of time. Our findings are consistent with previous, local or regional-scale studies, which generally found negative effects of human land use, and positive effects of landscape semi-natural habitat on bumblebee diversity (*5*, *13*, *27*). By analysing a collation of data spanning two continents, we show the generality of these effects, and also highlight the importance of landscape land-use history in shaping responses. Importantly, our findings show that responses to land use interact strongly with climate change and with the position of populations within species’ climatic niches.

Interestingly, bumblebee occupancy is reduced most by human land use in the centre of species’ temperature niches. A previous study of vertebrate species showed that populations are more likely to be lost in human-disturbed land uses near the edge of species’ thermal niches, specifically where populations are near their upper realized temperature limit (*44*). At the same time, a study of insects showed that species affiliated with cooler and wetter conditions were more likely to be absent in human land uses than those affiliated with warmer and drier conditions (*40*). These previous results are expected, given that human-disturbed land uses have hotter and drier local climates than natural habitats (*59*, *60*). Our finding of disproportionate losses in human land uses in the centre of species’ temperature niches is not consistent with local climatic changes being the principal mechanism.

Climate warming, which moves populations closer to species’ upper thermal limits, reduced occupancy in natural habitats, but not consistently in human land uses. This contrasts somewhat with the results of a study on birds in North America, where declines in response to habitat loss were strongest where summer temperatures had warmed the most (*61*). The strong negative response of bumblebees to human land use in the temperature niche centre, and the negative effect of climate change in natural habitats, may be associated with factors such as resource limitation (*62*, *63*), rather than local climatic changes. At the same time, the relative resilience of bumblebee populations to land use at the upper edge of species’ thermal niche possibly points towards some local adaptation to climatic conditions, as suggested by previous studies (*64*). On the other hand, other studies have suggested limited local adaptation to climate in bumblebees (*36*), which is consistent with the disproportionate declines of bumblebee occupancy seen at species’ upper thermal limits in response to recent climate change (*3*). Thus, it would appear that the way interactive effects of climate change and land-use change play out across the niches of bumblebee species is more complicated than appears to be the case for other groups of species.

Surprisingly, land-use intensification, measured here in terms of pesticide and fertilizer application rates, had relatively little effect on bumblebee biodiversity. However, we caution that the available global data on application of agricultural chemicals are resolved only at a coarse spatial resolution of 5 arc-minutes (approximately 10 km at the Equator) (*58*, *65*). Similar studies to ours have also found weak effects of fertilizer application rate using the same dataset (*41*), while a study of UK bees found detectable, but weak, effects of pesticide (specifically neonicotinoid) exposure on population trends (*20*). The results of these analyses of field data, using coarse-scale estimates of pesticide application rates, contrast strongly with results based on experimental exposure of bees to neonicotinoid pesticides both in the lab and the field (*19*, *23*). There is thus an urgent need for more detailed and finely resolved continental or global spatial data on the application of specific pesticides, as well as biodiversity data that can be associated more precisely with these application data.

Our results highlight the growing impacts on biodiversity from interactions among pressures, which will likely intensify rapidly in the coming decades. Based on the current credible policies and actions of the world’s countries, the global average climate is expected to warm by around 2.7^∘^C by 2100 (*66*). This will likely lead to decreases, on average, in bumblebee diversity (*3*, *14*, *37*), with our results highlighting particularly important declines in natural habitats. While, in general, Europe and North America are expected to see a reduction in cropland area in future, with relatively little change in agricultural intensity compared to other world regions (*67*), ongoing changes in agricultural practices are highly likely. We highlight the complex interplay of land-use and climatic factors in shaping bumblebee biodiversity, which need to be considered when predicting changes in biodiversity. Our findings also point to some potential interventions that could mitigate losses of bumblebee biodiversity. In particular, efforts to return areas of agriculture to a natural state, and to restore patches of natural habitat within farmed landscapes, are likely strongly to benefit bumblebees, and probably also other insect groups (*41*). However, any efforts to improve bumblebee biodiversity through changes in land use are unlikely to be successful unless the rate of climate change is also reduced.

Using our models to infer historical changes in bumblebee occupancy suggests that modification of landscapes, combined with recent climate change, has reduced bumblebee biodiversity within natural habitats almost as much as in human-modified areas, creating a ‘shifted baseline’ (*57*) in the present day. While agricultural areas experience the direct impacts of habitat loss, many natural habitats are exposed to degradation of their surrounding landscapes, while both natural and human-modified areas are exposed to the effects of climate change. Land-use impacts are often quantified using spatial analyses, comparing sampled biodiversity between natural and modified areas (*68*). The fact that natural habitats are composed of depauperate communities, heavily impacted by changes in their landscapes, means that such estimates are likely to underestimate land-use impacts substantially (*69*).

Understanding changes in biodiversity through time using a spatial database of species records is inevitably subject to certain limitations. Most importantly, such an analysis does not permit a consideration of the dynamics of biodiversity change (*70*). However, time-series data for insects are lacking in most regions, making a spatial analysis the only way to infer insect biodiversity changes at continental to global scales. Another limitation, mentioned earlier, is the reliance of large-scale studies on coarsely resolved estimates of the pressures on biodiversity. While the availability of fine-scale land-use/land-cover data has improved rapidly in recent years (*71*), we still lack fine-scale estimates of the application of agricultural chemicals, a particularly important pressure on flying insect biodiversity (*10*). To estimate the position of populations within species’ temperature niches, we use information on species’ realized distributions with respect to temperature. Such realized temperature limits may not correspond with the fundamental physiological limits of species, but estimates of the latter (*72*) are available for too few species to be useful in large-scale analyses such as ours. A previous study showed that realized thermal limits are clearly associated with occupancy change in response to recent climate change (*3*), suggesting that our approach is valid. Finally, all correlative analyses are limited in their ability to deduce causal mechanisms with certainty. The complexity of the interactive effects we document highlights a need for more experimental studies or detailed field-level assessments to investigate the mechanistic basis for the patterns we report.

In conclusion, we show that interactions between climate change and land use drive rapid and substantial changes in bumblebee biodiversity across both natural habitats and human-modified land. Given the importance of bumblebees for pollinating wild plants (*73*) and agricultural crops (notably apples, pears, melons, peppers, cucumbers, tomatoes, and many nuts and soft fruits (*54*)), these changes are likely to have important effects on both natural ecosystems and on our ability to grow food. Mitigating further losses of bumblebee diversity, and potentially reversing past declines, will require strong mitigation of climate change combined with restoration of natural habitats. To predict future changes in biodiversity, including of bumblebees, we must account for the complex interaction of climate and land-use pressures that are currently affecting species.

## Materials and Methods

### Data on Responses to Land Use

Data describing the occurrence of bumblebees in different land-use types were derived from the PREDICTS database (*56*). The PREDICTS database collates biodiversity records from individual studies that conducted snapshot spatial samples of biodiversity along land-use or land-use-intensity gradients (*55*). Studies included in the database had to meet a number of criteria: 1) that the study, or at least its methods, were published; 2) that biodiversity was sampled at more than one location; 3) that the study considered the effect of human activities on a set of sampled taxa, with the level of human activity varying among sampled locations; and 4) that the sampling protocol was the same across sampled sites. Any records without geographical coordinates were omitted. In this study, we focused on the occurrence (i.e., simple presence or absence) of taxa that were identified to the species level.

The final dataset that we used in this study were derived from 52 original studies, and consist of 17,555 records for 49 bumblebee species, collected across 1,593 sites in 13 different countries in North America, and Western, Northern and Southern Europe: Canada, United States, Belgium, Estonia, France, Germany, Ireland, Italy, Netherlands, Serbia, Sweden, Switzerland and United Kingdom (Figure 6). The original samples of bumblebees that were collated in the PREDICTS database were collected in the field between 2000 and 2011.

**Figure 6:**
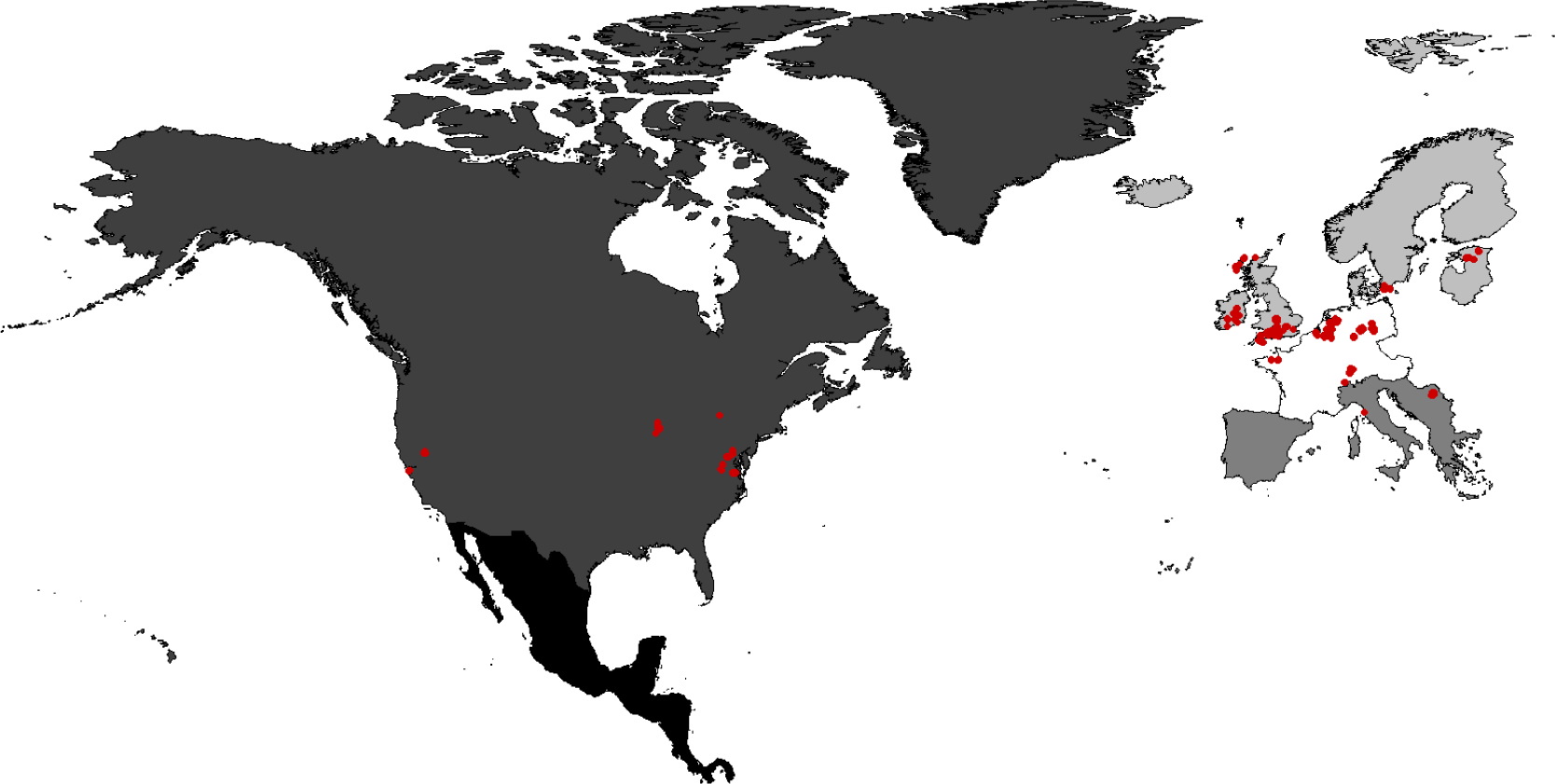
Location of sites used in the analysis. Base map shows the UN sub-regions from which bumblebee data were obtained: Northern America, Central America, Northern Europe, Western Europe and Southern Europe. Note that Mexico is classed within Central America in the UN sub-region scheme. Map is plotted with geographical coordinates using the WGS 1984 datum

### Explanatory Variables

We considered land use in our analysis by dividing sampled sites into either natural habitats or human-modified land. In the PREDICTS database, land use is classified into one of six broad categories: primary vegetation (natural habitat never having been recorded as being destroyed), secondary vegetation (habitat recovering to its natural state after being destroyed in the past by human actions or extreme natural events), plantation forests (areas used to cultivate woody crops), cropland (areas used to cultivate herbaceous crops, including as fodder for livestock), pasture (areas used regularly or permanently to graze livestock), and urban (areas of human settlement or other buildings, or areas managed for human recreation). We treated primary and secondary vegetation as natural habitats, and all other land-use types as human-modified.

We investigated how variables describing surrounding habitat conditions, land-use intensity (specifically, chemical application) and species’ temperature niche properties shape the occupancy of bumblebees in both natural and human-modified habitats.

Information on habitat condition in the landscapes surrounding sampled sites comprised estimates of the availability of natural habitat, as well as the length of time that landscape habitat has been substantially modified by humans. Availability of surrounding natural habitats was measured as the percentage of natural habitat in the 5 × 5-km grid cell within which the biodiversity sample was taken. Original mapped estimates of different land-use types were from a down-scaled land-use projection for 2005 (the mid-point of original field sampling of data included in this analysis; see previous sub-section) at 30-arc-second spatial resolution (approximately 1 km at the Equator) (*74*). These land-use maps estimate the fraction of each grid cell covered by five out of the six land-use types recognised in the PREDICTS database: primary and secondary vegetation, cropland, pasture and urban. Following our classification of overall land-use type (see above), we summed the primary and secondary vegetation maps to obtain estimates of natural habitat. We projected and resampled this map of the fractional cover of natural habitat at 30-arc-second resolution to an equal-area grid (Behrmann projection) at 1-km resolution, using bilinear interpolation (implemented using the projectRaster function of the raster R package Version 3.5-29; (*75*)). We then aggregated the resulting 1-km map by a factor of five, calculating the mean fractional cover of natural habitat for each 25-cell spatial cluster, yielding the final map at 5-km resolution. We derived estimates of the duration of substantial human habitat modification from a land-use reconstruction at 0.5° spatial resolution. Specifically, we used the reconstruction from the harmonized land-use estimates created for the fifth assessment report of the Intergovernmental Panel on Climate Change (*76*). The original historical land-use estimates used in this harmonization were from the HYDE model Version 3.1 (*77*). The harmonized historical land-use estimates describe reconstructed changes in the fractional cover of primary vegetation, secondary vegetation, croplands, pastures and urban areas from 1500 to 2005. We measured the duration of substantial human habitat modification as the number of years since a 0.5° grid cell was first 30% converted to human-modified land-use types (croplands, pastures or urban areas). Grid cells that have never yet reached the conversion threshold of 30% are considered to have a duration of substantial landscape modification of 0 years. A 30% loss of natural habitats from landscapes is a level above which the most sensitive species are thought to be substantially negatively impacted (*78*).

We considered the density (kg/ha) of application of two classes of chemicals: pesticides and fertilisers. We derived recent modelled estimates of pesticide application density across a 5-arc-minutes (approximately 10 km at the Equator) global spatial grid (*58*). We used the low rather than high modelled estimates from this dataset, to be conservative, but tested the robustness of our models to using the high estimates instead. We obtained estimates of fertiliser application rates from global modelled estimates, also at 5-arc-minutes spatial resolution (*65*). Estimates are available of total nitrogen, phosphorous and potassium application rates on each of 17 major global crops. We summed application rates for all three nutrients and all 17 crops to generate an estimate of total fertiliser application rate.

For information on species temperature niche properties, we used previously published methods and associated records of bumblebee occupancy (*3*). Specifically, we estimated: 1) the exposure of a species’ population to extreme high temperatures relative to its realized temperature niche in a baseline period before the onset of rapid climate changes (1901 — 1975); and 2) the change in exposure caused by climate change between the baseline period and a more recent period (2000 — 2014). Realized temperature niche limits were estimated for each species as the mean of the five lowest minimum monthly temperatures and the maximum of the five highest maximum monthly temperatures across all bumblebee spatial records in the baseline period. To measure the exposure of each species’ population to extreme high temperatures in either the baseline or recent period, we then derived the maximum monthly temperatures for all twelve months in each baseline or recent year. These maximum monthly temperatures were rescaled such that a value of 0 equates to the minimum realized temperature niche limit of the species, and a value of 1 to the maximum niche limit. We then averaged the 12 monthly values to derive an annual mean exposure. Finally, we averaged the annual exposure estimates across all years in either the baseline or recent period. Thus, we were able to estimate, for any biodiversity record, the average exposure to extreme high temperatures in the baseline period, or the change in exposure between the baseline and recent periods. Monthly minimum and maximum temperature estimates for the period 1901 to 2015 were obtained from the CRU global gridded climate reconstruction Version 3.24.01 (*79*), which was the most recent version of this dataset available at the time the bumblebee climatic niche properties were estimated.

### Statistical Analysis

To analyse the presence or absence of bumblebee species at the sampled locations, we used binomial generalised linear mixed-effects models, to account for the hierarchical structure of the PREDICTS database. In all models, we used a binary response variable representing the presence or absence of each species at each sampled location. Species were considered absent if they were not recorded but had been targeted by a particular study (typically these species were sampled in at least one of the sites within each study). As random effects, we fitted: 1) Study identity, to control for differences in sampling protocols and broad geographic differences; 2) Site identity nested within study identity, to control for site-specific factors; and 3) Species identity, to account for variation in responses among species unrelated to variation in the environmental driver variables. Although there may be important differences between the two continents in average occupancy of bumblebees, as well as in responses to the considered explanatory variables, the size of the dataset and the fact that sampling was much less intensive in North America than Europe prevented the inclusion of such variation in the models.

We considered a candidate set of models with different fixed-effects structures. First, we fitted a null, intercept-only model. Second, a model with only land use as a fixed effect. Third, models fitting, as well as land use, all possible combinations of the different broad classes of explanatory variables (landscape habitat, land-use intensity, and climatic niche properties), and also interactions between these additional explanatory variables and land use (Table 1). We assessed model fit to the data using AIC values. For each model *i* in the set of models *I*, we calculated the difference in its AIC compared to the best-fitting model, Δ_*i*_, and the model AIC weight, *AIC*_*w*_:

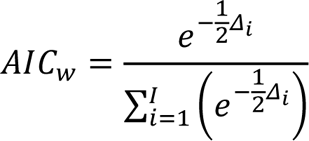

We also calculated the sum of AIC weights of all models containing a particular class of explanatory variable. In addition to comparing the statistical support for models by comparing AIC values and AIC weights, we also estimated model explanatory power by computing pseudo R^2^ values following (*80*), conducted model diagnostic tests using the DHARMa package Version 0.4.5 in R (*81*), and tested for spatial autocorrelation in the residuals associated with each underlying study using Moran’s test, implemented in the spdep package Version 1.2-8.

To reveal the impacts that environmental changes to date have had on bumblebee biodiversity across both natural habitats and human-modified habitats, we used our final set of models to infer biodiversity changes across the sampled sites, based on the values of the model explanatory variables at those sites at the time of biodiversity sampling (or as close as possible - see above). These estimates thus included the impacts on both natural and modified habitats of climate change, and the modification of landscapes by human land use. Estimates of biodiversity at the time of sampling were expressed as a percentage difference compared to a baseline condition in which we assumed that landscapes were entirely composed of natural habitat, thus with no history of human modification, subject to no application of fertilizers and pesticides, and with no change in climate experienced. To derive model-estimated differences between baseline and contemporary conditions, we randomly generated 1,000 estimates of percentage difference to represent both model and parameter uncertainty. Thus, each estimate was generated by: 1) drawing at random from among the candidate models, weighted by their AIC weights (see above); 2) drawing a set of parameters from this chosen model based on a multivariate normal distribution, with means being the model-estimated coefficients, the standard deviation from the coefficient uncertainties, and parameter correlations derived from the variance-covariance matrix; 3) using these parameters to estimate average bumblebee probability of occurrence under baseline and contemporary environmental conditions; and 4) calculating the percentage difference in average occurrence probability between baseline and contemporary conditions.

## Acknowledgments

### Funding

Royal Society International Exchanges grant to TN and JK (IE161031)

UK Natural Environment Research Council grant to TN (NE/R010811/1)

Royal Society University Research Fellowship to TN (UF150526)

### Author Contributions

Conceptualization: TN, JK, PS and JJW

Formal analysis: TN

Funding acquisition: TN and JK

Methodology: TN, JK, PS and JJW

Writing - original draft: TN

Writing - review & editing: JK, PS and JJW

## Model Diagnostic Checks

**Figure S1:**
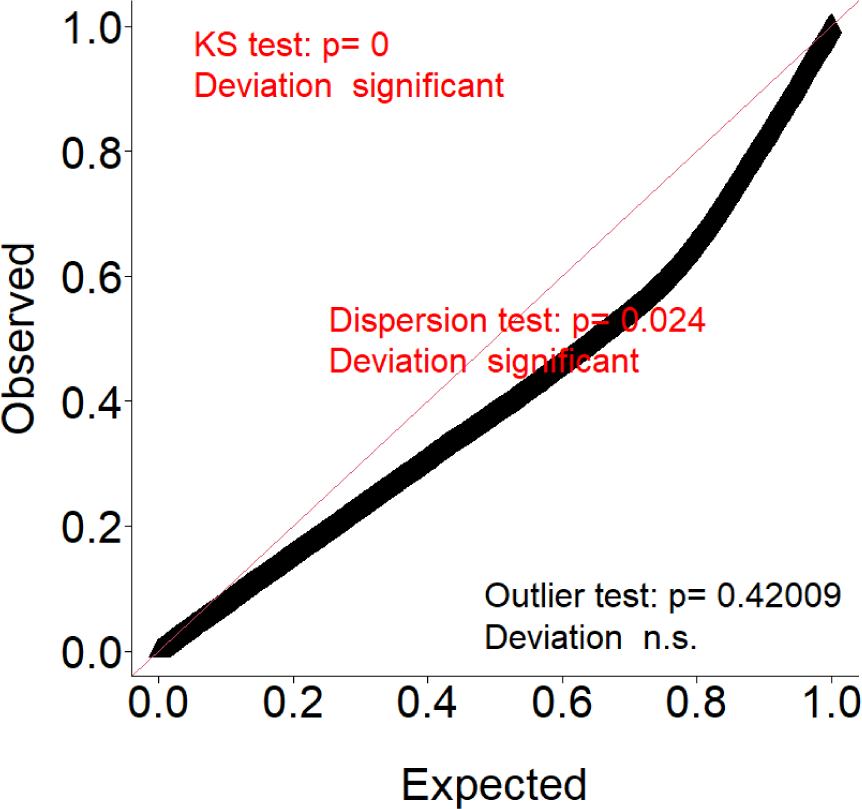
Q-Q plot of scaled model residuals compared to the expected distribution of residuals. This plot was produced directly from the DHARMa R package Version 0.4.6

**Figure S2:**
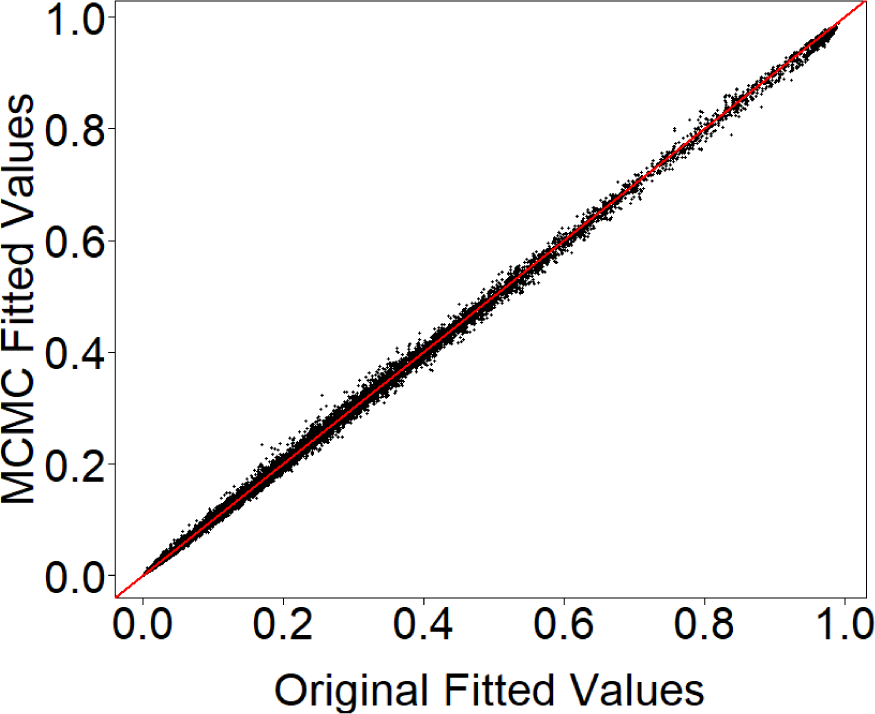
Comparison of fitted values from the main best-fitting model and a model with identical structure fitted using MCMC. This comparison was made to check for robustness of model-estimated probabailities of presence, given the violation of the assumptions of standard parametric statistical tests, shown in Figure S1. The MCMC model was fit using the MCMCglmm R package Version 2.34

**Figure S3:**
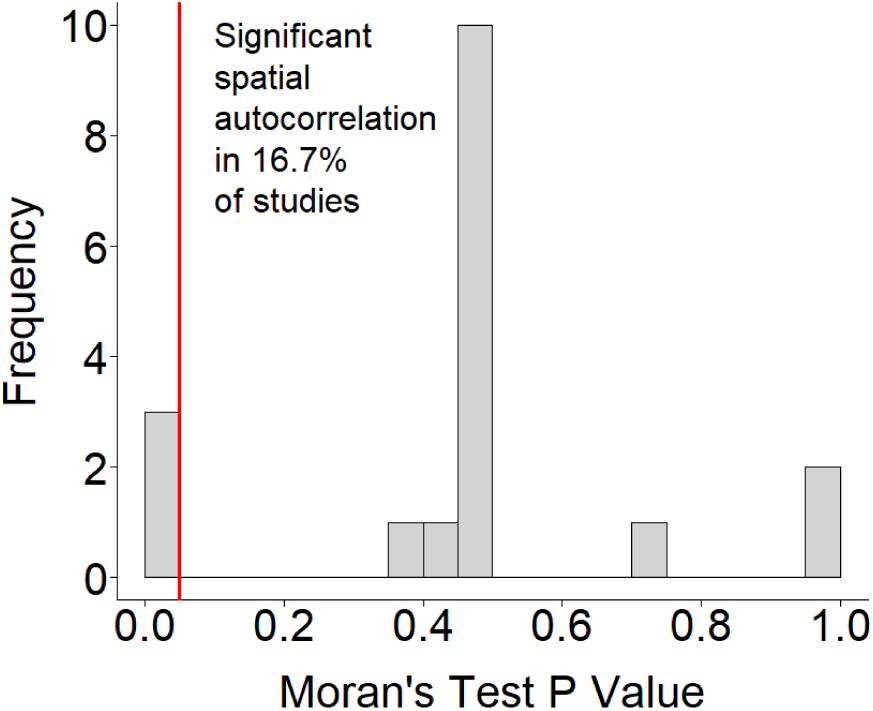
Distribution of P values from a series of Moran’s tests for spatial autocorrelation in the residuals associated with each individual study. For each study, we calculated the average residual for each sampled site, and then ran a Moran’s test for spatial autocorrelation in these average residuals. Moran’s tests were performed using the spdep R package Version 1.2-8

